# TOMATO FERTIGATION MANAGEMENT UNDERPINS A PHYSIOLOGICAL AND MOLECULAR REPROGRAMMING UNDER SALT STRESS

**DOI:** 10.64898/2026.10.20.760371

**Authors:** José A. Martín-Coll, Sara Gómez-Jiménez, Raúl López-Blanco, Albano Carrera, Carlos García-Saez, Mónica Calvo-Polanco, Pablo Albertos, Oscar Lorenzo

**Affiliations:** Dept. of Botany and Plant Physiology, Excellence Research Unit of Agricultural Production and Environment (AGRIENVIRONMENT), Institute for Agribiotechnology Research (CIALE), Faculty of Biology, University of Salamanca. C/ Río Duero 12, 37185 Salamanca, Spain; BISITE Research Group, University of Salamanca, C/ Espejo 2, 37007 Salamanca, Spain; Nutricontrol S.L. Polígono Industrial Cabezo Beaza, C/ Bucarest 26, 30395 Cartagena, Región de Murcia, Spain

**Keywords:** Tomato, transcriptome, soil salinity, nitrate efficiency, stress combination

## Abstract

Agricultural crops are often exposed to concurrent environmental stresses, with soil salinization and intensive fertilization representing two major threats to sustainable food production. Here, we investigated the effect of nitrogen availability on salinity tolerance in tomato (*Solanum lycopersicum*) through integrated physiological, agronomic, and transcriptomic analyses. Our results confirm that salinity-induced declines in chlorophyll content were mitigated by moderate nitrate reduction, while vegetative development remained largely unaffected. Likewise, reproductive traits under moderate nitrate supply were comparable to those observed in fully fertilized plants. Interestingly, fruit quality traits in terms of sugars and lycopene content, were enhanced particularly under salinity combined with reduced nitrate supply. Consistent with these physiological responses, transcriptomic analyses across the different nitrate regimes revealed extensive salinity-induced transcriptional reprogramming, with the largest response observed under moderate nitrate supply. Together, these findings highlight the potential of precise fertilization management to improve both productivity and fruit quality in saline environments.

**SIGNIFICANCE:** Nitrogen management represents a critical yet underexplored determinant of crop performance under salinity. Moderate nitrate reduction sustained tomato growth, reproductive development, and fruit quality while eliciting the strongest transcriptional response to salt stress. In contrast, severe nitrate limitation was associated with molecular reprogramming linked to translational processes and resource allocation toward fruit quality traits. These results identify nitrate availability as a key regulator of salinity acclimation and highlight optimized nitrogen management as a promising strategy to improve nitrogen-use efficiency in salt-affected cropping systems.

## INTRODUCTION

Modern agriculture faces the formidable challenge of increasing global production to meet the food demands of an increasing population within the adverse context of climate change (Aishwarya & Kumar, 2024). Factors such as erratic precipitation, extreme heatwaves, soil salinization, and nutrient deficiencies—particularly nitrate limitation— constitute critical abiotic stresses that directly impair crop development and constrain global yields (Fahad et al., 2021). Elucidating the physiological, molecular, and genetic mechanisms of adaptation is fundamental to implementing sustainable and resilient agricultural practices capable of enhancing both crop productivity and fruit quality under a changing climate.

The present study focuses on tomato (*Solanum lycopersicum*), a globally ubiquitous crop and one of the most widely consumed vegetables, with an estimated annual intake of 61,9-198,9 kg per capita (Quinet et al., 2019). Beyond its economic importance, tomato plays a vital role in human nutrition as a rich source of bioactive compounds, including essential minerals, dietary fiber, vitamins (notably ascorbic acid), and potent antioxidants such as lycopene (Ali et al., 2020).

Despite its adaptability to diverse environments, tomato production is significantly compromised by environmental pressures. While abiotic stress research is frequently conducted under controlled conditions focusing on isolated factors, plants in natural ecosystems typically encounter a combination of simultaneous environmental constraints (Rivero et al., 2014a). In response, plants deploy integrated and synchronized physiological, metabolic, and molecular mechanisms to achieve acclimation. Understanding these specific regulatory networks is essential for developing advanced genetic improvement strategies and enhancing environmental resilience.

Among essential elements, nitrogen (N) is a critical macronutrient for plant growth and a primary constituent of amino acids, chlorophyll, and nucleic acids (Qin et al., 2019; Xu et al., 2012). Tissue N concentration serves as a robust indicator of nutritional status; as a core component of the chlorophyll molecule, its deficiency becomes a primary limiting factor for achieving optimal yields (Mu & Chen, 2021).

Previous research has demonstrated that both nitrogen deficiency and excess limit productive performance. Núñez-Ramírez et al. (2017) investigated this effect in tomato plants subjected to various fertilization rates (250, 500, 750, and 1000 kg N/ha), observing that intermediate concentrations yielded the highest productivity. While excessive nitrogen promotes vegetative biomass at the expense of reproductive development, N restriction impairs overall growth and subsequent reproductive organ formation (Núñez-Ramírez et al., 2017). Notably, that study found no significant differences in fruit quality parameters, such as total soluble solids (TSS), titratable acidity (TA), or the TSS/TA ratio. Conversely, a review by Dorais et al. (2000) indicates that high N fertilization negatively impacts fruit colour, maturation time, and uniformity, while reducing TSS content. This phenomenon is attributed to a “dilution effect,” where rapid fruit growth and increased water accumulation dilute primary metabolites and sugars. Conversely, moderate nitrate limitation may slightly enhance sugar concentration, provided that photosynthetic activity is not compromised (Dorais et al., 2000).

In modern agriculture, N deficiency is typically managed through synthetic fertilizers; however, precision in dosage is paramount. Optimizing N application is necessary not only for yield and cost-efficiency but also for environmental sustainability, preventing issues such as water eutrophication and soil salinization. Inorganic fertilizer consumption reached 185 million tonnes of nutrients in 2022, with nitrogen accounting for 58%—a 37% increase since 2000 (“World Food and Agriculture – Statistical Yearbook 2024,” 2024). This intensive use, driven by the quest for higher productivity, is intrinsically linked to anthropogenic soil salinization.

Currently, approximately 1,381 million hectares (Mha) globally (10.7% of the Earth’s land surface) are affected by salinity or sodicity. While present on all continents, over two-thirds of these areas are located in arid and semi-arid regions, with the largest affected areas in Australia, Argentina, and Kazakhstan, among others. Future projections suggest that global aridity trends could increase the salt-affected land surface to 24–32% by the end of the century (Park et al., 2018). Salinization already compromises 10% of both irrigated and rainfed croplands, threatening global food security. While causes can be natural (e.g., weathering, seawater intrusion), anthropogenic factors such as poor irrigation management and inappropriate fertilizer use significantly exacerbate the problem (“Global Status of Salt-Affected Soils,” 2024)

Regarding salinity stress, mounting evidence suggests that plants actively repress growth as an adaptive strategy to maximize survival (H. Zhang et al., 2022). This highly sensitive response program is activated even under mild stress through priming mechanisms. In the initial stages, stress signaling pathways actively suppress anabolic activities and vegetative growth, often before cellular energy status is compromised (Praveen et al., 2023). In tomato, salinity stress significantly reduces relative water content, restricts growth, and inhibits photosynthesis. Furthermore, reproductive development is impaired by ionic imbalances in floral tissues (P. Zhang et al., 2016). The accumulation of toxic Na^+^ and Cl^−^ ions in female floral tissues (style and ovaries) disrupts reproductive timing by delaying anthesis, while increasing flower abortion, reducing fertility, and impairing seed set in tomato crops(Ghanem et al., 2009). Similarly, in *Arabidopsis*, salinity has been shown to disrupt gametophyte development, resulting in non-viable seeds (Cackett et al., 2022).

To mitigate these adverse effects, plants deploy complex responses involving the accumulation of osmoprotectants (e.g., proline and soluble sugars) and the up-regulation of enzymatic and non-enzymatic antioxidants. In tomato specifically, salt stress has been observed to enhance fruit organoleptic quality (P. Zhang et al., 2016). Controlled salinity levels can significantly improve qualitative parameters by promoting the accumulation of lycopene, starch, sucrose, and amino acids (Tang et al., 2020; P. Zhang et al., 2016).

## METHODS

### Plant material and growth conditions

For the experimental trials, a tomato variety (*Solanum lycopersicum*) provided by Meridiem Seeds was utilized. This variety exhibits an indeterminate growth habit. In accordance with standard commercial greenhouse management practices for such varieties, apical pruning was performed once the plants reached the sixth inflorescence. Additionally, lateral shoots were regularly removed to prevent excessive vegetative growth and to promote fruit development.

Seeds were sown in plastic seedling trays filled with vermiculite within the greenhouse. During the seedling stage, plants were fertigated twice weekly with a commercial NPK formulation (Fertiplant). Once the plants had 3 o 4 true leaves were transplanted into coconut coir bags (Projar), an inert substrate chosen for its optimal water-holding capacity and porosity.

Experiments were conducted in a specialized greenhouse module equipped with three independent drainage benches, each dedicated to a specific nitrate fertilization level. Environmental parameters were strictly regulated using an automated climate control system integrated with heating and evaporative cooling units. High-precision sensors continuously monitored temperature and relative humidity (RH), maintaining a thermal regime of 22°C/18°C (day/night). Supplemental lighting was provided to ensure a 16:8 h (light:dark) photoperiod, while retractable shading screens were utilized to modulate excessive solar radiation. Each bench featured an autonomous fertigation system. Nutrient solutions were prepared in individual tanks with three distinct nitrate formulations: 100%, 50%, and 25% of the standard concentration. The nutrient solution was daily mixed using a timer-controlled agitator (mixer tashia 125 L 12 V cc s.N°: 231191 – 231192) to ensure chemical homogeneity and prevent salt precipitation. A diaphragm dosing pump (Type RF 409.2-50e, sera GmbH, Immenhausen, Germany) delivered the concentrated nutrient solution into the irrigation circuit for dilution with potable water. Downstream from the mixing point, the pipeline split into two distinct branches: one for control plants and the other for plants subjected to salt stress. This experimental design enabled the simultaneous evaluation of six conditions (3 nitrate levels × 2 treatments: control and salinity). Three tomato plants were grown per bag, with eight bags allocated per condition, resulting in a total of 24 plants per treatment and 48 plants per drainage bench. The experimental setup comprised the following six conditions: 100% nitrate, 100% nitrate + 200 mM NaCl, 50% nitrate, 50% nitrate + 200 mM NaCl, 25% nitrate, and 25% nitrate + 200 mM NaCl. This design enabled the evaluation of tomato responses to stress combination, integrating salinity stress with contrasting levels of nitrate availability, ranging from optimal nutrition (100%) to severe nitrate limitation (25%).

The fertilization protocol was based on a modified Hoagland solution, specifically formulated such that nitrogen was provided through a single salt source. This experimental design facilitated precise modulation of N concentrations across the three tested levels by adjusting only one component of the nutrient mix, thereby ensuring that the concentrations of all other macro- and micronutrients remained constant. The final chemical composition and elemental concentrations in the irrigation water post-dilution were as follows: boric acid (H_3_BO_3_) 2,86 mg/L, calcium chloride, anhydrous (CaCl_2_) 554,9 mg/L, copper II chloride, anhydrous (CuCl_2_) 0,045 mg/L, EDTA, iron sodium salt (C_10_H_12_N_2_NaFeO_8_.3H_2_O) 33 mg/L, magnesium sulphate, anhydrous (MgSO_4_) 240, 325 mg/L, manganese chloride, tetrahydrate (MnCl_2_ 4H_2_O) 1,81 mg/L, molybdic acid sodium salt, dihydrate (Na_2_MoO_4_ .2H_2_O) 0,025 mg/L, potassium chloride (KCl) 372,7 mg/L, potassium phosphate, monobasic, anhydrous (KH_2_PO_4_) 136,025 mg/L, zinc chloride, anhydrous (ZnCl_2_), 0,11 mg/L y ammonium nitrate (NH_4_NO_3_) 480,24 mg/L (100%).

### Stress treatment

Salinity was induced through the manual addition of NaCl to the nutrient solution. To ensure homogeneity in water delivery, each automated irrigation cycle performed for the control groups was manually replicated for the stress groups, providing an identical volume of solution at a final concentration of 200 mM NaCl.

### Hydroponic system setup and fertigation control

Irrigation timing and frequency were regulated via an automated solenoid valve coupled to the main water supply. The injection rate of the concentrated nutrient solution was regulated by a variable frequency drive (Eura drives), which controlled the operation of the metering pump. To ensure a stable flow rate, a pressure regulator was installed at the water main outlet, maintaining a constant operating pressure of 2.5 bar.

The nutrient solution dilution ratio was determined by the pump’s operating frequency. To establish the precise mixing percentage, both the total system outflow (delivered via 48 emitters) and the pump’s specific contribution were empirically measured across the entire range of operating frequencies. For the latter, a calibration curve was constructed by plotting the pump’s rotational speed (Hz) against its injection flow rate (L/h). Given that the total system flow remains constant under the regulated pressure, the fertilizer mixing ratio was calculated based on the specific volume injected by the pump into the irrigation circuit.

Each irrigation line featured two monitoring nodes for substrate moisture, temperature, and electrical conductivity (EC). These parameters were recorded and monitored remotely, allowing for real time adjustments to maintain the target experimental conditions for each trial.

### Determination of sugar content by HPLC

Once the tomatoes reached optimal ripeness, they were harvested for physicochemical analysis. Five biological replicates were established per experimental condition (100C_1, 100C_2, 100C_3, 100C_4, and 100C_5), each consisting of a randomized pool of five tomatoes. To preserve the physicochemical and organoleptic characteristics, the tomatoes were processed on the same day of harvest.

For each replicate, five tomatoes were homogenized using a blender until a uniform liquid extract was obtained. Aliquots of 50 mL were collected and stored at –20 °C for subsequent extraction and quantification of sugars and lycopene.

Glucose and fructose were quantified by High-Performance Liquid Chromatography (HPLC) using an Agilent 1200 HPLC system equipped with a refractive index detector (RID). A Rezex™ ROA-Organic Acid H+ column (300 x 7.8 mm i.d.) was utilized for the separation. The eluent was 0.005 N sulfuric acid, delivered isocratically at a flow rate of 0.5 mL min^-1^, with the column temperature maintained at 60°C.

### Determination of lycopene content

Lycopene determination was performed using a Thermo Vanquish UHPLC system coupled to a Thermo Orbitrap Q Exactive Focus high-resolution mass spectrometer. Chromatographic separation was achieved on an Ascentis Express C18 column (100 x 2.1 mm i.d., 2.7 µm particle size). The mobile phase consisted of 100% acetonitrile maintained at a flow rate of 2 mL min^-1^.

Ionization was conducted using an electrospray ionization (ESI) source operating in fast polarity switching mode (positive/negative). Mass spectra were acquired in full-scan mode ranging from m/z 100 to 1000 at a resolution of 70,000 FWHM.

### Total RNA extraction

Leaf samples were harvested at two points in time after the induction of salt stress: at the onset of the first and the third inflorescence anthesis. For each experimental condition, three biological replicates were established, consisting of tissue pools from plants located in identical relative positions within the cultivation bags (positions 1 to 3). Samples were snap-frozen in liquid nitrogen and stored at –80 °C for subsequent analysis.

For RNA-Seq analysis, samples corresponding to the second sampling event performed 8 weeks (58 days) post-stress induction were selected. Leaf tissue was mechanically homogenized, and total RNA was extracted and purified from 50 mg of sample using the E.Z.N.A.® Plant RNA Kit (Ref. R6827). RNA concentration was determined via NanoDrop spectrophotometry. Furthermore, the integrity and quality of the genetic material were assessed using the RNA ScreenTape Assay (Ref. 5067-5576) on a TapeStation 4150 system (S/N DEDAB00634). For sequencing, 20 µL aliquots per sample were prepared at an approximate concentration of 1000 ng/µL, ensuring an RNA Integrity Number (RIN) > 6 and 28S/18S ratios > 1. Sequencing services were outsourced to BGI Tech Solutions (Poland).

### RNA sequencing and differential expression analysis

Three to four samples per experimental condition were prepared by diluting 1 µg of total RNA in nuclease-free water to a final concentration of 50 ng/µL and sent to Beijing Genomics Institute (BGI, Hong Kong, China; https://www.bgi.com). RNA concentration and integrity were assessed at BGI facilities using a Bioanalyzer. Based on RNA integrity number (RIN) values and 28S/18S rRNA ratios, the three highest-quality samples per condition were selected for library preparation and sequencing. Strand-specific cDNA libraries were prepared using the DNBSEQ Eukaryotic Strand-specific Transcriptome Resequencing workflow and sequenced using DNA nanoball technology (DNBSEQ™), generating 150-bp paired-end reads with approximately 30 million clean reads per sample. Sequencing read quality was assessed using FastQC version 0.11.9 (Andrews, 2010). Reads were aligned to the *Solanum lycopersicum* ITAG4.1 reference genome using HISAT2 version 2.1.0 (Kim et al., 2019). Transcript abundance was estimated using StringTie version 2.2.1 guided by the ITAG4.1 gene annotation. Gene-level read counts were extracted using the prepDE.py script provided with StringTie.

Differential gene expression analysis was performed in R version 4.4.2 using DESeq2 version 1.46.0 (Love et al., 2014). Genes with at least 10 reads in a minimum of three samples were retained for analysis. A generalized linear model based on the negative binomial distribution was fitted using a factorial design including nitrate concentration (25, 50, and 100), stress treatment (Control and NaCl), and their interaction (∼ Nitrate per Stress). The nitrate concentration of 25 and the Control treatment were used as reference levels. The transcriptional response to NaCl was evaluated separately at each nitrate concentration by estimating the corresponding NaCl versus Control contrasts from the fitted model. Genes with a Benjamini–Hochberg-adjusted *p*-value ≤ 0.05 were considered DEG, without imposing an additional log_2_ fold chang (log_2_FC) threshold.

To identify genes whose transcriptional response to NaCl depended significantly on nitrate concentration, a likelihood ratio test (LRT) was performed by comparing the full interaction model (∼ Nitrate per Stress) with a reduced additive model (∼ Nitrate + Stress). Genes with an adjusted *p*-value ≤ 0.05 were considered to exhibit a significant nitrate per stress interaction. DEGS were further classified as up-regulated, down-regulated, or non-significant according to the direction of their log_2_FC and adjusted p-value. Overlaps and regulatory transitions among nitrate concentrations were examined to identify consistently regulated and nitrate-dependent responses.

Principal component analysis (PCA) was performed using variance-stabilized count data obtained with the DESeq2 variance-stabilizing transformation. To compare the magnitude of the NaCl response across nitrate concentrations, absolute log_2_FC values were examined for genes consistently up- or down-regulated across all three concentrations and for genes exhibiting a significant nitrate per stress interaction. Differences among nitrate concentrations were assessed using Friedman tests, followed by paired Wilcoxon signed-rank tests with Benjamini–Hochberg correction for multiple comparisons.

### GO enrichment

Gene Ontology (GO) enrichment analysis was performed using the clusterProfiler R package (Yu et al., 2012) and GO annotations obtained from the *Solanum lycopersicum* ITAG4.1 genome annotation. Gene-to-GO associations were extracted from the annotation dataset and used to construct a custom annotation database for enrichment analysis. Only genes with at least one GO annotation were included in the enrichment analysis. Accordingly, the background universe consisted of genes retained after DESeq2 expression filtering that had at least one GO annotation (10,104 genes).

DEGs from the NaCl versus Control comparisons at nitrate concentrations of 25, 50, and 100 were classified as up- or down-regulated according to their log_2_FC direction and a Benjamini–Hochberg-adjusted *p*-value ≤ 0.05. GO enrichment was assessed separately for up- and down-regulated genes at each nitrate concentration, as well as for genes consistently up- or down-regulated across all three concentrations, considering only GO-annotated genes within each set. Comparative enrichment analysis was performed using the enricher function within the compare Cluster framework, with a custom gene-to-GO mapping derived from ITAG4.1 annotations.

Additionally, GO enrichment was performed on GO-annotated genes showing a significant nitrate per stress interaction, as identified by the likelihood ratio test. Enrichment significance was assessed using the Benjamini–Hochberg procedure, and GO terms with adjusted p-values ≤ 0.05 were considered significantly enriched. Significantly enriched terms were visualized using dot plots displaying fold enrichment, gene counts, and adjusted p-values.

### Statistical analysis

All statistical analyses were performed using R Statistical Software (v4.2.1; R Core Team 2023)(R Core Team, 2023). Rosnet test, from EnvStats package(Millard, 2013), was used to identify outliers. Car(Fox et al., 2024), and agricolae (de Mendiburu, 2023) and rstatix (Kassambara, 2023b) packages were used for multiple comparisons analysis, where ANOVA or Kruscall Wallis test for nonparametric data were assessed significant differences and Duncan test to find significantly different samples. Ggplot2(Wickham, 2016) was used to graph the data and ggpubr (Kassambara, 2023a) to add statistics to the graphics. The figures have been exported using the officer package in R(Gohel et al., 2024).

## RESULTS

### Reduced nitrate supply preserves chlorophyll accumulation but impairs vegetative growth under saline conditions

During the 15-week vegetative growth evaluation period, plant height increased steadily in all treatments, although growth rates were strongly affected by both nitrate availability and salinity (Figure 1A). Plants fertigated with 100% nitrate displayed the highest stem elongation, whereas progressive reductions in nitrate supply resulted in lower final plant height. Salt stress further restricted vegetative growth, particularly under the 25% nitrate treatment, indicating an additive effect of salinity and nitrogen limitation on shoot development.

**Figure 1.**
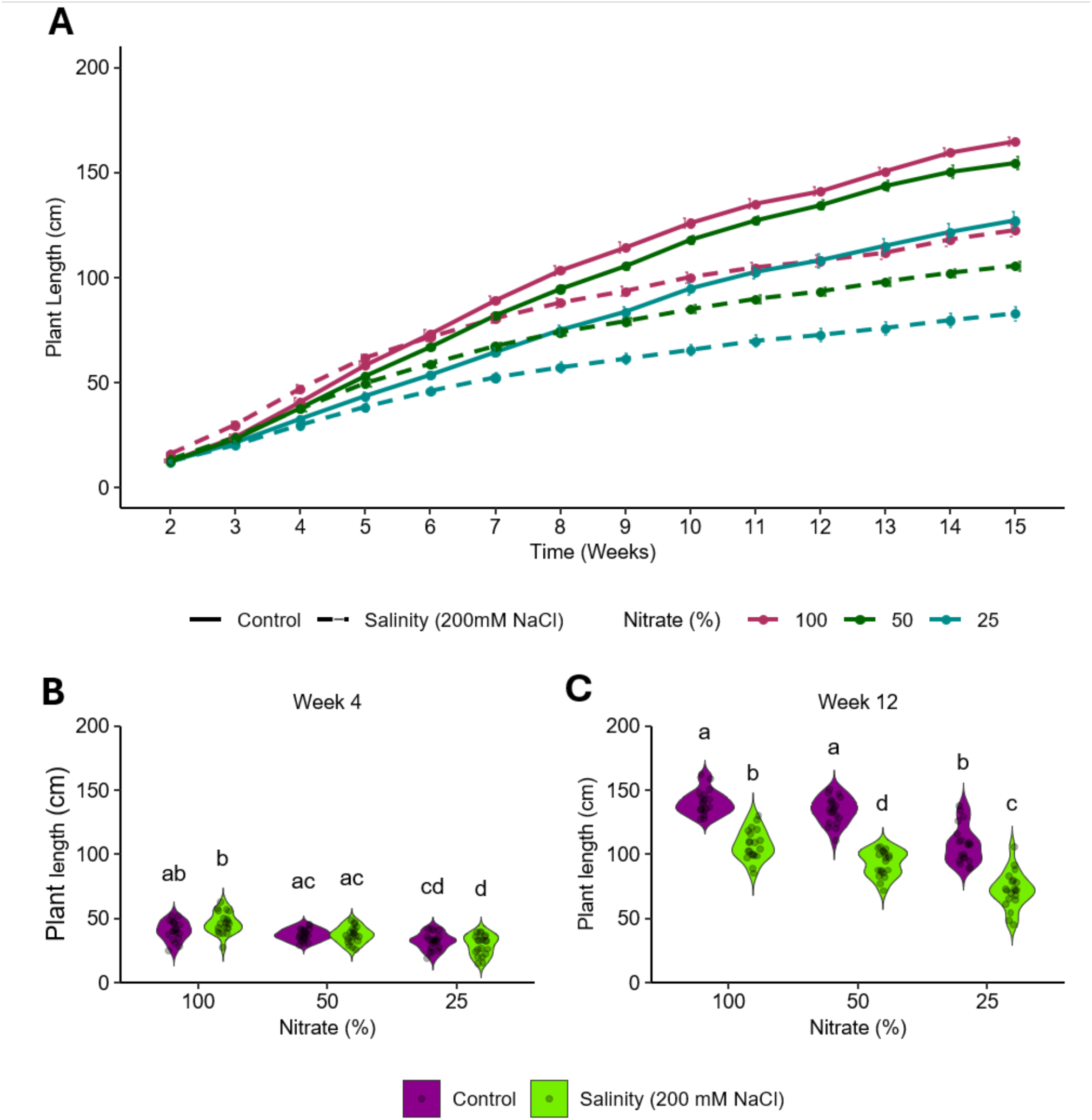
Weekly growth dynamics of tomato plants grown under different nitrogen fertigation regimes combined with control and saline stress conditions. (A) Plant height development, expressed as the mean height of the 24 plants within each treatment, from week 2 (following acclimation to coconut coir and the onset of salinity treatment) to week 15 (apical meristem removal in the 100% nitrate control treatment). (B) and (C) show the distribution of plant height within each treatment at weeks 4 and 12, respectively. Data were analysed using an Aligned Rank Transform ANOVA (ART), followed by pairwise comparisons with Bonferroni adjustment. Different letters indicate significant differences among treatments (*p* < 0.05).

Under non-saline conditions, plants supplied with 100% nitrate exhibited the greatest vegetative growth, reaching approximately 165 cm at week 15, followed by plants receiving 50% nitrate (155 cm). In contrast, plants grown with 25% nitrate showed a marked reduction in stem elongation, reaching final heights close to 125 cm. These differences became increasingly evident throughout plant development and were already statistically significant at week 12 (Figure 1C).

Salinity significantly reduced plant height regardless of nitrate supply. Plants exposed to 200 mM NaCl and fertigated with 100% nitrate reached approximately 122 cm at week 15, whereas those receiving 50% nitrate attained around 105 cm. The greatest growth inhibition was observed under the combined treatment of salinity and severe nitrate restriction (25%), where plant height remained below 85 cm. At week 12, control plants supplied with 100% and 50% nitrate showed significantly greater stem elongation than all salt-stressed treatments, while the lowest values were recorded in plants receiving 25% nitrate under saline conditions (Figure 1C).

Leaf chlorophyll content, estimated through SPAD measurements, increased progressively throughout plant development in all treatments (Figure 2A). Nevertheless, chlorophyll accumulation was strongly influenced by nitrate availability. Under control conditions, plants supplied with 100% nitrate consistently exhibited the highest SPAD values, reaching approximately 67 units by week 15. Reducing nitrate availability caused a gradual decline in chlorophyll content, with plants receiving 25% nitrate showing the lowest values during the later stages of development.

**Figure 2.**
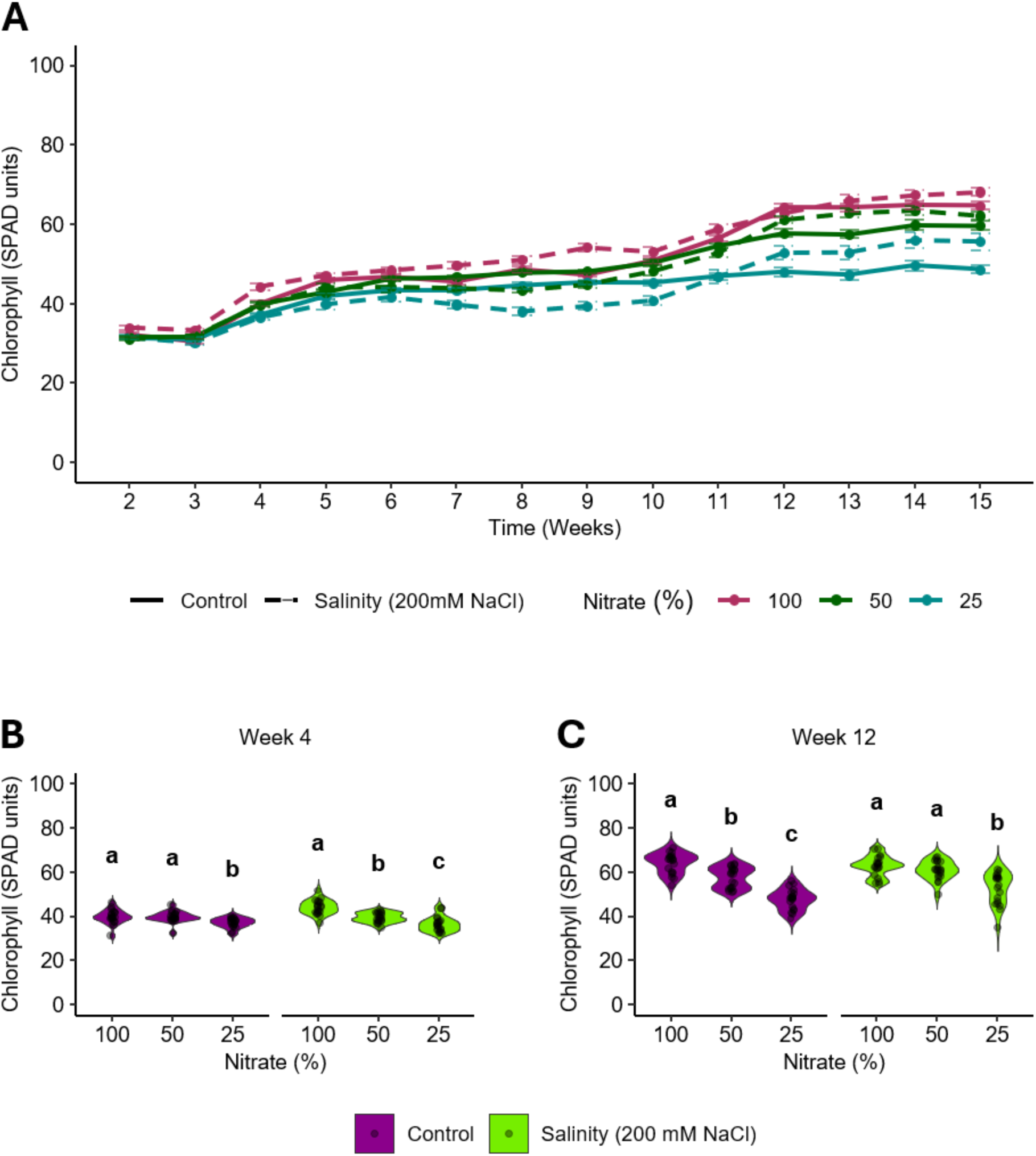
Weekly chlorophyll dynamics in tomato plants grown under different nitrogen fertigation regimes combined with control and saline stress conditions. **(A)** Mean chlorophyll content of the 24 plants within each treatment from week 2 to week 15. **(B)** and **(C)** show the distribution of chlorophyll content within each treatment at weeks 4 and 12, respectively. Data were analysed using an Aligned Rank Transform ANOVA (ART), followed by pairwise comparisons with Bonferroni adjustment. Different letters indicate significant differences (*p* < 0.05) among nitrate levels within each treatment condition (control and saline stress).

Differences in chlorophyll status were already apparent at week 4, when plants supplied with 25% nitrate exhibited significantly lower SPAD values than those receiving 50% or 100% nitrate under both control and saline conditions (Figure 2B). By week 12, the effect of nitrate nutrition became even more pronounced, with chlorophyll content decreasing progressively with nitrate reduction irrespective of salinity treatment (Figure 2C). Notably, plants supplied with 50% nitrate maintained SPAD values comparable to those observed under full nitrate nutrition, suggesting that moderate nitrate restriction did not substantially compromise leaf chlorophyll accumulation.

Overall, nitrate availability exerted a stronger effect on leaf chlorophyll content than salinity, whereas both factors markedly influenced vegetative growth. Notably, plants receiving 50% nitrate-maintained growth and chlorophyll levels relatively close to those of plants supplied with 100% nitrate, while severe nitrate restriction (25%) significantly reduced both parameters under control and saline conditions.

### Nitrate availability determines reproductive precocity under saline conditions

To characterize the effects of nitrate availability and salinity on reproductive development, flowering and fruit set dynamics were analysed using a Cox proportional hazards model. This approach allowed the inclusion of censored observations corresponding to plants that had not completed a given phenological stage by the end of the experimental period.

The opening dynamics of the first six inflorescences revealed a clear effect of nitrate availability on reproductive precocity (Figure 3A). Under control conditions, plants supplied with 100% and 50% nitrate showed very similar flowering patterns, with rapid progression through successive inflorescences and high proportions of flowering plants. In contrast, plants grown under 25% nitrate exhibited a consistent delay in anthesis, particularly from the third inflorescence onwards, indicating that severe nitrate restriction slowed the transition between reproductive stages.

**Figure 3.**
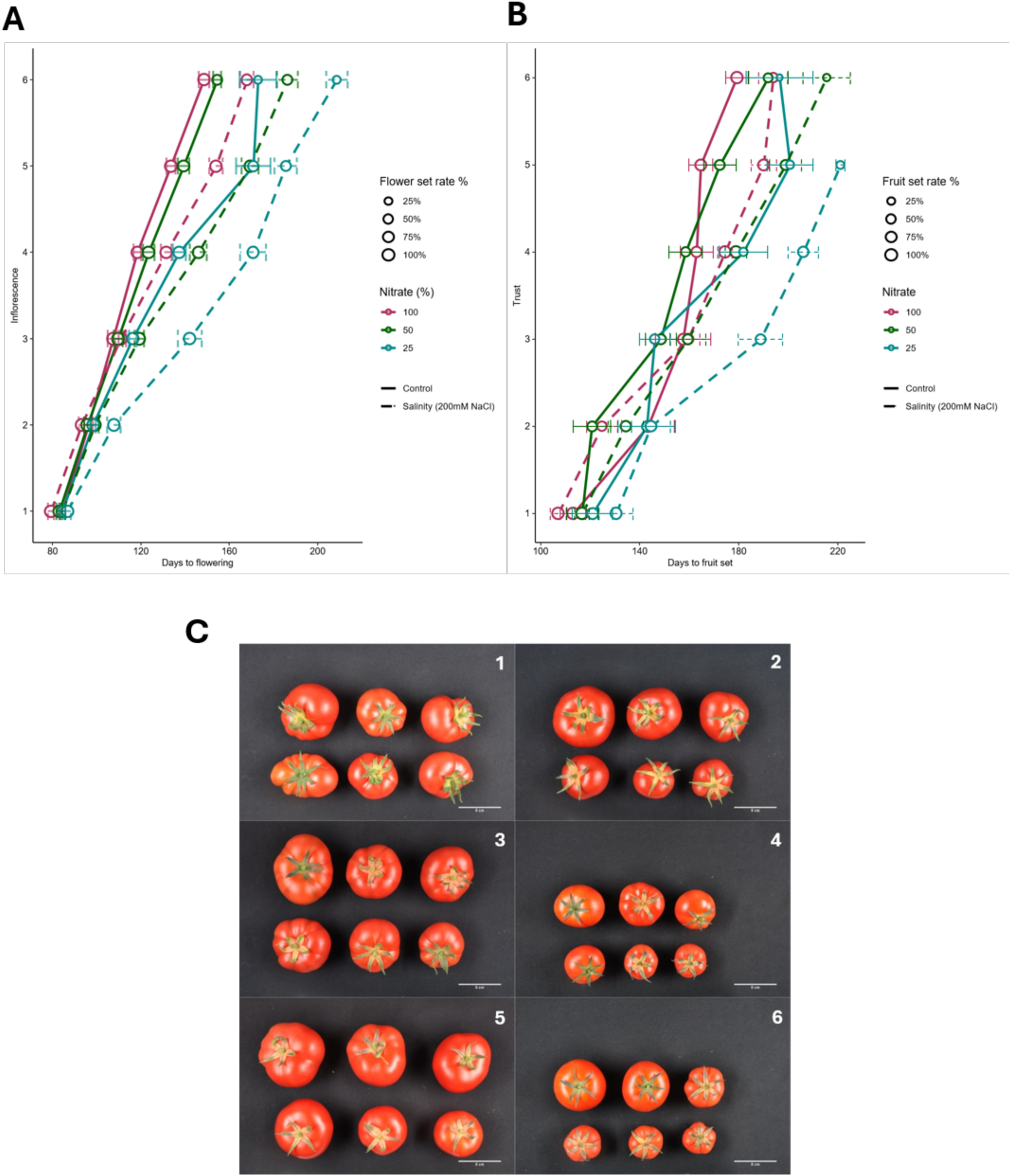
Reproductive phenology across six successive inflorescences and representative fruit development in tomato plants grown under different nitrogen fertigation regimes combined with control and saline stress conditions. **(A)** Mean time to flowering and **(B)** fruit set for each inflorescence in tomato plants grown under different nitrogen fertigation regimes combined with control and saline stress conditions. Point size represents the percentage of plants (out of 24) that successfully set at least one fruit within each truss. **(C)** Representative tomato fruit harvest illustrating the variability in fruit size, morphology and colour among the different experimental groups: **(1)** 100% nitrate + control, **(2)** 100% nitrate + salinity stress, **(3)** 50% nitrate + control, **(4)** 50% nitrate + salinity stress, **(5)** 25% nitrate + control and **(6)** 25% nitrate + salinity stress.

Salt stress generally delayed flowering compared with the corresponding controls. However, nitrate availability strongly modulated this response. Plants supplied with 100% nitrate under salinity maintained flowering kinetics similar to those observed in the control treatments and consistently outperformed plants grown with 25% nitrate. Delays associated with the 25% nitrate treatment became progressively greater with increasing inflorescence order, suggesting cumulative effects of nitrogen limitation on reproductive development.

A similar pattern was observed for fruit set dynamics (Figure 3B). Treatments receiving 100% nitrate reached fruit set earlier than plants supplied with reduced nitrate concentrations, regardless of salinity conditions. Under saline conditions, high nitrate availability partially mitigated the delay in fruit establishment, whereas plants exposed to the combined treatment of salinity and 25% nitrate showed the slowest progression through reproductive stages. This treatment also displayed lower percentages of plants reaching fruit set in the upper trusses before the end of the experiment.

Overall, reproductive phenology was more strongly affected by nitrate availability than by salinity alone. Moderate nitrate reduction (50%) maintained flowering and fruit set dynamics comparable to those observed under full fertilization, whereas severe nitrate limitation delayed both anthesis and fruit establishment throughout the crop cycle.

### Salinity preserves fruit sugar accumulation and lycopene content independently of nitrate availability

Glucose and fructose concentrations exhibited remarkably similar response patterns across treatments (Figure 4A, 4B). Under control conditions, soluble sugar accumulation showed a clear dependence on nitrate availability. Fruits from plants supplied with 100% and 50% nitrate displayed comparable glucose and fructose concentrations, whereas severe nitrate restriction (25%) significantly reduced the accumulation of both carbohydrates. In this treatment, glucose and fructose concentrations declined to values below 3%, resulting in significant differences with respect to the 100% nitrate treatment. In contrast, salinity substantially modified this response. Under 200 mM NaCl, the negative effect of nitrate limitation on sugar accumulation disappeared and a reverse trend was observed. Fruits from salt-stressed plants maintained high concentrations of both glucose and fructose across all nitrate treatments, with the highest values generally recorded under 25% nitrate supply. Consequently, the reduction in soluble sugar content observed under severe nitrate restriction in control plants was not detected under saline conditions.

**Figure 4.**
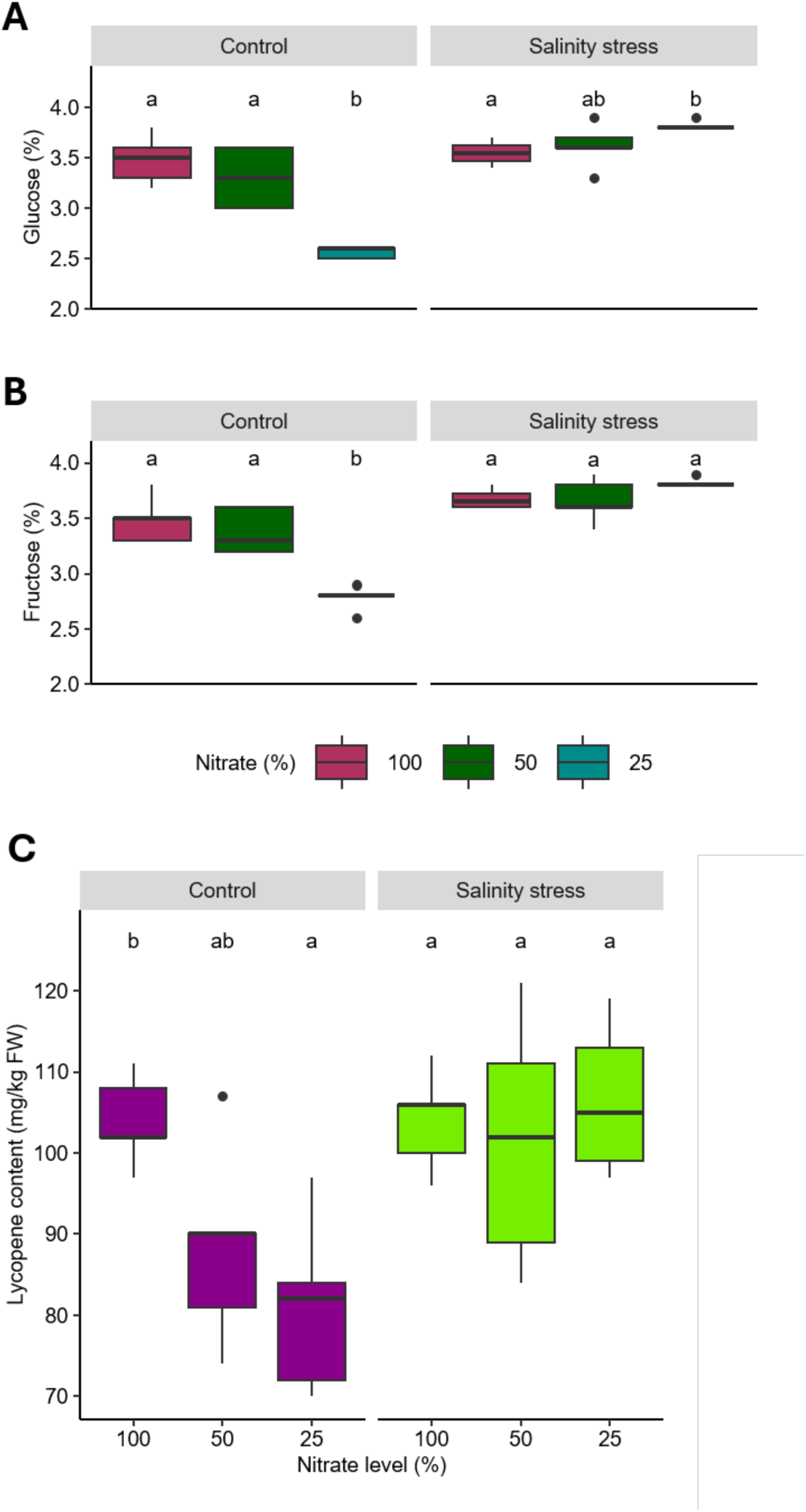
Distribution of glucose (A), fructose (B), and lycopene (C) concentrations in tomato fruits harvested from plants grown under different nitrate fertigation regimes under control and saline stress conditions. For each treatment, metabolite concentrations were quantified from five biological replicates. Data were analysed using an Aligned Rank Transform ANOVA (ART), followed by pairwise comparisons with Bonferroni adjustment. Different letters indicate significant differences (*p* < 0.05) among nitrate levels within each treatment condition (control and saline stress).

These results indicate that while nitrogen deficiency limits carbohydrate accumulation under non-stress conditions, salinity promotes the maintenance of elevated glucose and fructose concentrations independently of nitrate availability.

Lycopene accumulation was strongly influenced by nitrate availability under control conditions (Figure 4C). Fruits harvested from plants supplied with 100% nitrate exhibited the highest lycopene concentrations, whereas a progressive reduction was observed as nitrate availability decreased. Significant differences were detected between the 100% and 25% nitrate treatment, indicating that severe nitrate restriction negatively affected carotenoid accumulation during fruit ripening.

In contrast, salinity substantially modified the lycopene response to nitrate availability. Under 200 mM NaCl, no significant differences were detected among nitrate treatments, and lycopene concentrations remained consistently high across all nutritional regimes. Notably, fruits from salt-stressed plants receiving 50% and 25% nitrate accumulated lycopene levels equal to or higher than those measured in fruits from the 100% nitrate control treatment.

Therefore, while nitrate availability regulated lycopene accumulation under control conditions, salinity prevented the reduction in lycopene concentration associated with nitrogen deficiency and maintained elevated carotenoid levels independently of nitrate supply.

### Nitrate availability modulates the extent and magnitude of the transcriptional response to salt stress in tomato

To investigate how nitrate availability influences the transcriptional response to salt stress, RNA-seq analysis was performed in tomato plants grown under three nitrate concentrations (25, 50, and 100) in the presence or absence of NaCl. Principal component analysis (PCA) revealed differences in global transcriptional profiles across experimental conditions, with the first two principal components explaining 60% and 22% of the total variance, respectively (Figure S1A). Samples grown at the lowest nitrate concentration (25) clustered relatively close to each other, regardless of NaCl treatment, whereas a clearer separation between control and salt-stressed samples was observed at nitrate concentrations of 50 and 100. Differential expression analysis identified 2,704 (1236 Up + 1468 Down), 6,373 (2906 Up + 3467 Down), and 5,285 (2565 Up + 2720 Down) DEGs in response to NaCl at nitrate concentrations of 25, 50, and 100, respectively (Figure S1B). Thus, the largest number of NaCl-responsive genes was detected at 50, followed by 100 and 25. At all three concentrations, both up- and down-regulated genes were identified in relatively similar proportions, with a modest predominance of down-regulated genes (Fig 1B, C).

To determine the extent to which the NaCl response was conserved across nitrate concentrations, the overlap among up- and down-regulated DEG sets was examined using an UpSet plot (Figure 5A). A total of 1,314 genes were differentially expressed at all three nitrate concentrations (449 consistently up-regulated, 860 consistently down-regulated, and five showing opposite regulatory directions between concentrations). In addition, 637, 2,166, and 1,567 DEGs were exclusively detected at nitrate concentrations of 25, 50, and 100, respectively, considering both up- and down-regulated genes. Overall, the direction of the transcriptional response was largely maintained across nitrate concentrations. When considering genes differentially expressed in at least two nitrate conditions, only 34 genes showed opposite regulatory responses in at least one pairwise comparison. This predominance of consistent regulatory patterns was also illustrated by the alluvial plot showing transitions between up-regulated, down-regulated, and non-significant states across nitrate concentrations (Figure S1D).

**Figure 5.**
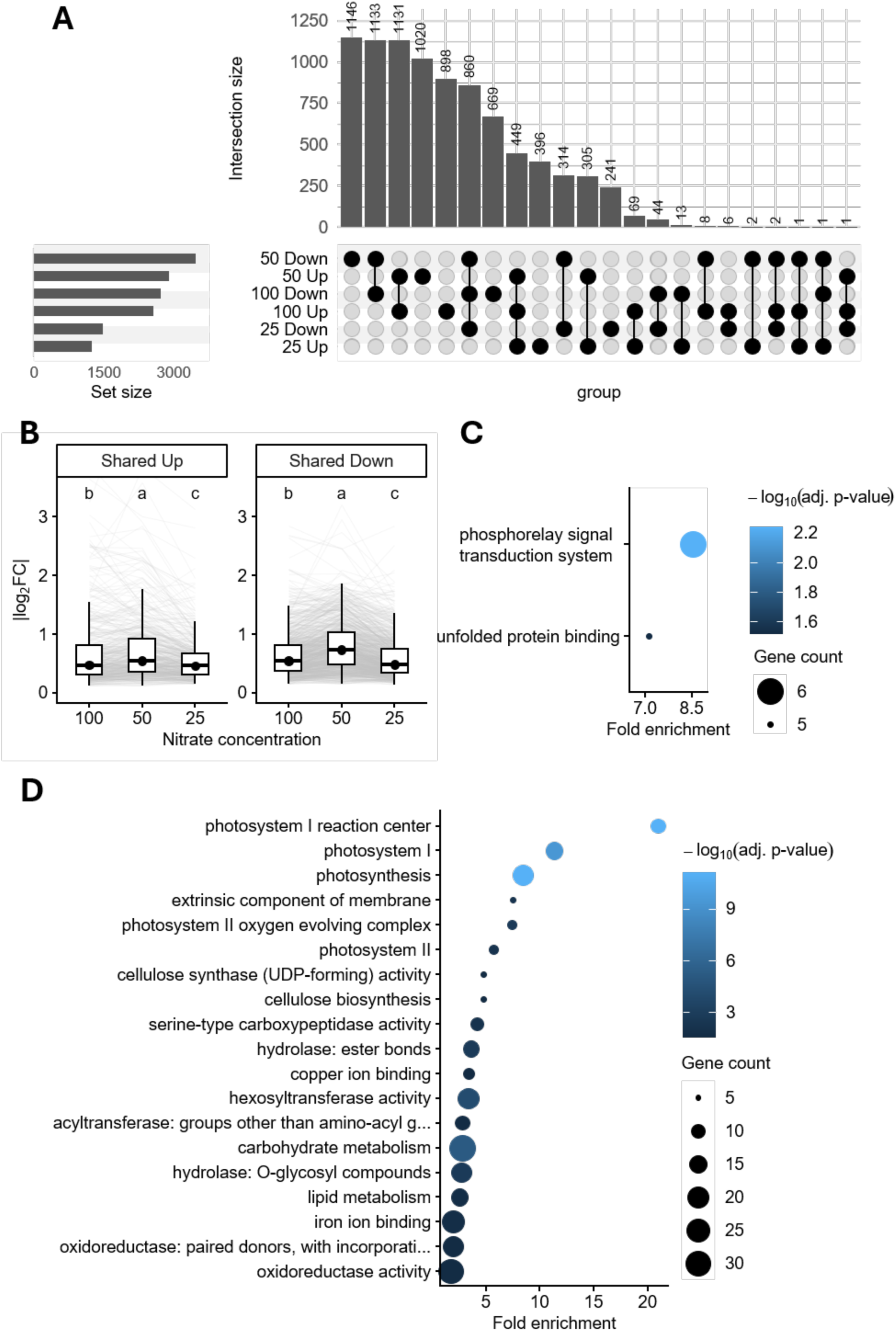
Shared transcriptional responses to NaCl across nitrate concentrations and their functional enrichment. **(A) Overlap of NaCl-responsive DEGs across nitrate concentrations.** UpSet plot showing the overlap among up- and down-regulated genes identified in the NaCl versus Control contrasts at nitrate concentrations of 25, 50, and 100. Genes were considered DEGs using an adjusted *p*-value ≤ 0.05. Horizontal bars indicate the total number of genes in each DEG set, whereas vertical bars indicate the size of each intersection. Filled circles connected by vertical lines indicate the DEG sets contributing to each intersection. **(B) Magnitude of the NaCl transcriptional response in genes consistently regulated across nitrate concentrations.** Boxplots showing the absolute log_2_FC (|log_2_FC|) of genes significantly up-regulated (Shared Up) or down-regulated (Shared Down) in the NaCl versus Control contrasts at nitrate concentrations of 100, 50, and 25. Grey lines connect values corresponding to the same gene across nitrate concentrations. Different letters indicate significant differences between nitrate concentrations according to paired Wilcoxon signed-rank tests with Benjamini–Hochberg correction, following a significant Friedman test. Genes were considered differentially expressed using an adjusted *p*-value ≤ 0.05. **(C) GO enrichment of NaCl-induced genes at different nitrate concentrations**. Dot plot showing significantly enriched Gene Ontology terms among genes up-regulated by NaCl at nitrate concentrations of 50 and 25. The x-axis indicates nitrate concentration. Point size represents fold enrichment relative to the GO-annotated background, and point colour represents the adjusted p-value. No significantly enriched GO terms were detected among up-regulated genes at nitrate concentration 100. Only terms with adjusted *p*-value ≤ 0.05 are shown. **(D) GO enrichment of NaCl-repressed genes at different nitrate concentrations.** Dot plot showing significantly enriched Gene Ontology terms among genes down-regulated by NaCl at nitrate concentrations of 100, 50, and 25. The x-axis indicates nitrate concentration. Point size represents fold enrichment relative to the GO-annotated background, and point colour represents the adjusted p-value. Only terms with adjusted *p*-value ≤ 0.05 are shown.

Although a substantial number of genes showed consistent regulation across nitrate concentrations, the magnitude of their response to NaCl differed. Comparison of absolute log_2_FC (|log_2_FC|) values revealed that both consistently up-regulated and consistently down-regulated genes exhibited the highest median response magnitude at nitrate concentration 50 (Figure 5B). For the 449 shared up-regulated genes, median |log_2_FC| values were 0.459, 0.545, and 0.471 at nitrate concentrations of 25, 50, and 100, respectively. A similar pattern was observed for the 860 shared down-regulated genes, with median values of 0.477, 0.731, and 0.540, respectively. Friedman tests followed by paired Wilcoxon signed-rank tests with Benjamini–Hochberg correction identified significant differences among nitrate concentrations in both groups (Figure 5B). Nevertheless, differences in median response magnitude were relatively modest, particularly among the shared up-regulated genes. These findings suggest that nitrate availability influences not only the number of NaCl-responsive genes but also the magnitude of the transcriptional response among genes consistently regulated across nitrate conditions.

To explore the functional characteristics of the conserved NaCl response, Gene Ontology (GO) enrichment analysis was performed separately for shared up- and down-regulated genes, considering only genes with available GO annotations. The shared up-regulated set showed significant enrichment for two GO terms, *phosphorelay signal transduction system* and *unfolded protein binding* (Figure 5C). In contrast, the shared down-regulated set exhibited a broader functional enrichment profile, including terms associated with photosynthesis and photosystem components, particularly photosystems I and II, as well as carbohydrate metabolism, cellulose biosynthesis, and several enzymatic activities related to hydrolysis and oxidation–reduction processes (Figure 5D).

GO enrichment was also examined separately for up- and down-regulated genes at each nitrate concentration. Among up-regulated genes (Figure S2A), the largest number of significantly enriched GO terms was detected at nitrate concentration 25, including categories associated with translation, ribosomal components, protein folding, and transcription. Only two significantly enriched terms were identified at 50, whereas no significant enrichment was detected at 100 (Figure S2A). In contrast, down-regulated genes showed a broader enrichment profile across nitrate concentrations, with the highest number of significant GO terms detected at 100, followed by 50 and 25. Notably, terms associated with photosynthesis and photosystem components were enriched among down-regulated genes at all three nitrate concentrations, whereas additional categories related to carbohydrate metabolism, cell wall-associated processes, and various enzymatic activities differed among conditions (Figure S2B).

Together, these results indicate that the transcriptional response to NaCl includes a substantial core of genes consistently regulated across nitrate concentrations, alongside differences in the number of responsive genes, the magnitude of their expression changes, and the functional enrichment patterns associated with each nitrate condition.

### A subset of genes exhibits a significant nitrate per salt stress interaction

To identify genes whose transcriptional response to NaCl was significantly influenced by nitrate concentration, a likelihood ratio test (LRT) was performed by comparing a full model including the nitrate per stress interaction with a reduced additive model. This analysis identified 287 genes showing a significant nitrate per stress interaction (adjusted *p*-value ≤ 0.05). Unlike the comparisons described above, which characterized the transcriptional response to NaCl at each nitrate concentration, this analysis identified genes for which inclusion of the interaction term significantly improved the model fit.

The transcriptional responses of these 287 genes were subsequently examined at each nitrate concentration (Figure 6A). The number of genes classified as non-significant (NS) increased from 78 at nitrate concentration 100 to 191 at 25, where only 65 and 31 genes were significantly up- and down-regulated, respectively. Thus, fewer genes within this interaction-associated subset exhibited significant transcriptional responses to NaCl at the lowest nitrate concentration.

**Figure 6.**
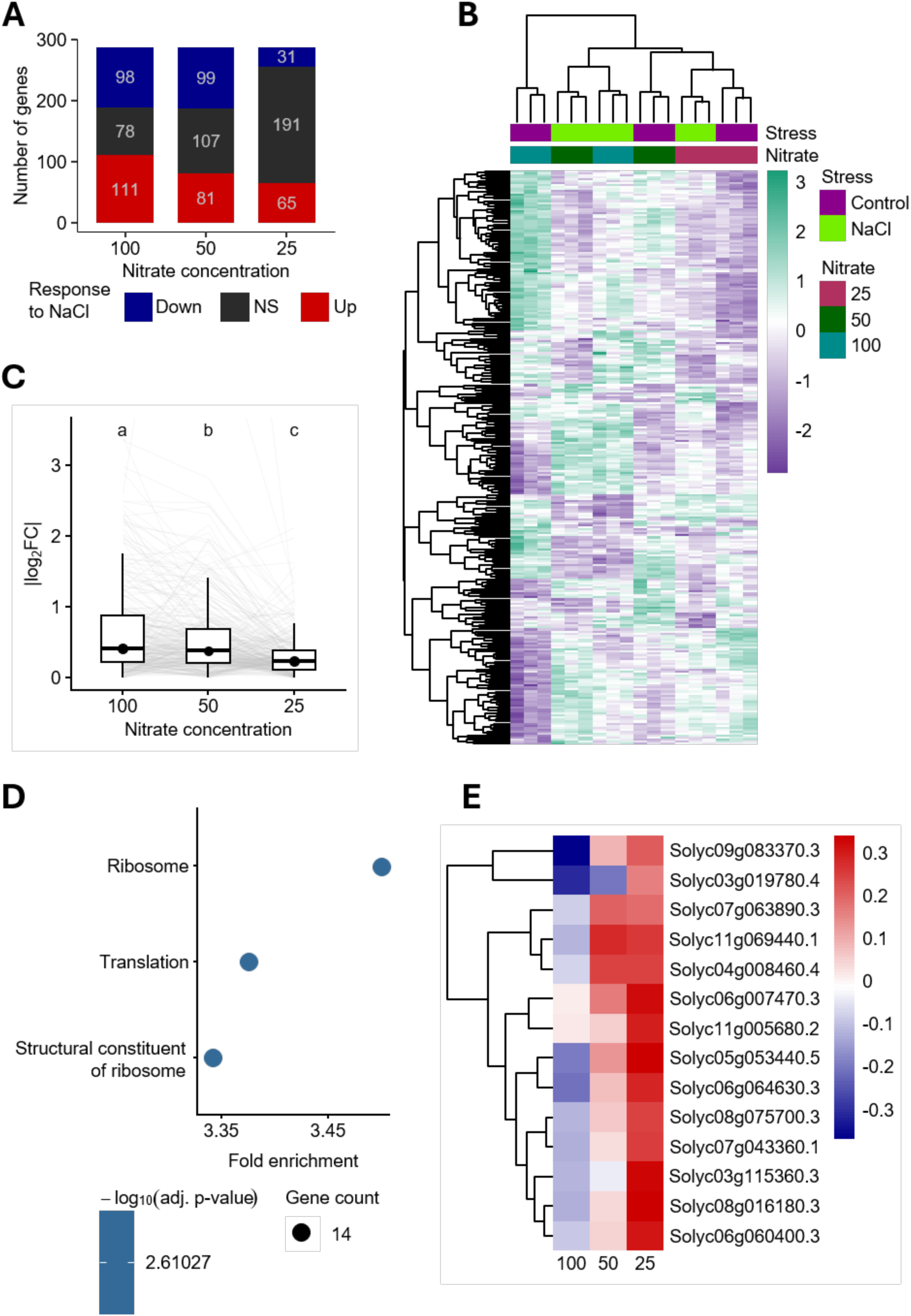
Transcriptional and functional characterization of genes showing a significant nitrate per salt stress interaction. **(A) NaCl-responsive genes among those showing a significant nitrate per stress interaction.** Stacked bar plot showing the number of up-regulated, down-regulated, and non-significant (NS) genes in the NaCl versus Control contrasts at nitrate concentrations of 100, 50, and 25, considering only the 287 genes identified by the likelihood ratio test as having a significant nitrate per stress interaction (padj ≤ 0.05). Genes were classified as up-regulated, down-regulated, or non-significant based on an adjusted *p*-value ≤ 0.05 in each NaCl versus Control contrast. Numbers within bars indicate the number of genes in each category. **(B) Expression patterns of genes showing a significant nitrate per stress interaction.** Heatmap showing the expression profiles of the 287 genes identified by the likelihood ratio test as having a significant nitrate per stress interaction (adjusted *p*-value ≤ 0.05). Variance-stabilizing transformed (VST) counts were scaled by gene as Z-scores to represent relative expression across samples. Rows and columns were hierarchically clustered, and sample annotations indicate nitrate concentration and NaCl treatment. Purple and green indicate relatively lower and higher expression, respectively. **(C) Magnitude of the NaCl response in genes showing a significant nitrate per stress interaction.** Boxplots showing the absolute log_2_FC (|log_2_FC|) of the 287 genes identified by the likelihood ratio test as having a significant nitrate per stress interaction (*padj* ≤ 0.05) at nitrate concentrations of 100, 50, and 25. Grey lines connect values corresponding to the same gene across nitrate concentrations. Different letters indicate significant differences according to paired Wilcoxon signed-rank tests with Benjamini– Hochberg correction, following a significant Friedman test. **(D) GO enrichment of genes showing a significant nitrate per stress interaction.** Dot plot showing the Gene Ontology terms significantly enriched among genes identified by the likelihood ratio test as having a significant nitrate per stress interaction. The x-axis represents fold enrichment relative to the GO-annotated background, point size indicates the number of genes associated with each term, and point colour represents the adjusted *p*-value. Only terms with adjusted *p*-value ≤ 0.05 are shown. **(E) NaCl response of genes contributing to significantly enriched GO terms.** Heatmap showing the log_2_FC of genes associated with the significantly enriched GO categories in the NaCl versus Control contrasts at nitrate concentrations of 100, 50, and 25. Rows represent individual genes and columns represent nitrate concentrations. Red and blue indicate positive and negative log_2_FC, respectively, whereas values close to zero are shown in white. Rows were hierarchically clustered according to similarity in response patterns.

Hierarchical clustering of variance-stabilized expression values for the 287 genes revealed heterogeneous expression patterns across experimental conditions (Figure 6B). Control samples at nitrate concentration 100 formed a distinct cluster, whereas control and NaCl-treated samples at nitrate concentration 25 clustered relatively close to each other. NaCl-treated samples at concentrations 50 and 100 also displayed relatively similar expression profiles.

The magnitude of the NaCl response was further evaluated by comparing absolute log_2_FC (|log_2_FC|) values among nitrate concentrations for the 287 genes (Figure 6C). The highest median |log_2_FC| was observed at nitrate concentration 100 (0.410), followed by 50 (0.379) and 25 (0.233). Friedman tests followed by paired Wilcoxon signed-rank tests with Benjamini–Hochberg correction revealed significant differences among all three nitrate concentrations. These results describe variation in response magnitude within the subset of genes identified as having a significant nitrate per stress interaction, with the lowest median response magnitude observed at nitrate concentration 25.

To investigate the functional characteristics of these genes, GO enrichment analysis was performed using the 165 interaction-associated genes with available GO annotations. Three significantly enriched terms were identified: *ribosome* (GO:0005840), *translation* (GO:0006412), and *structural constituent of ribosome* (GO:0003735), with fold enrichment values of approximately 3.3–3.5 and adjusted p-values < 0.05 (Figure 6D). All three terms were associated with the same 14 genes, indicating that the enrichment was driven by a small, overlapping group of genes related to ribosomal components and translation rather than three independent functional groups.

To further characterize their response to NaCl, the log_2_FC values of these 14 genes were examined across nitrate concentrations (Figure 6E). At nitrate concentration 25, all 14 genes exhibited positive log_2_FC values. At concentration 50, the log_2_FC values were generally lower but remained predominantly positive. Conversely, at nitrate concentration 100, log_2_FC values were predominantly negative. These results suggest a shift in the predominant direction of the NaCl response, from positive log_2_FC values at the lowest nitrate concentration towards predominantly negative values at the highest concentration.

Overall, the analysis revealed a subset of genes exhibiting a significant nitrate per salt stress interaction, characterized by differences in the magnitude and direction of their transcriptional responses across nitrate concentrations. Functional enrichment further identified a small group of ribosome- and translation-associated genes with nitrate-dependent expression patterns.

## DISCUSSION

Vegetative growth is typically one of the first physiological processes affected by salinity. Factors such as reductions in cell expansion, alterations in water relations and ion toxicity limit shoot development (Roșca et al., 2023). Exposure to saline conditions substantially reduced plant height across the vegetative phase, confirming the strong sensitivity of tomato growth to osmotic and ionic stress (Al-Gaadi et al., 2024). This salinity induced growth reductions have been widely associated with decreased water uptake, impaired nutrient acquisition and the metabolic costs required to maintain cellular homeostasis under high NaCl concentrations (Rivero et al., 2014b). Plant growth was also strongly influenced by nitrate availability. Under non saline conditions, plants receiving the full nitrate dose exhibited the greatest stem elongation, whereas growth progressively declined as nitrate supply was reduced. The marked reduction observed under the 25% nitrate treatment reinforces the importance of adequate nitrogen nutrition for sustaining leaf development, carbon assimilation and biomass accumulation (Paponov et al., 2021). The marked reduction in growth observed under the 25% nitrate treatment confirms that nitrogen availability became a major limiting factor for plant development, even in the absence of salinity. The similar growth observed in the 50% and 100% nitrate treatments suggests that moderate nitrate reduction did not compromise vegetative development, indicating a certain degree of flexibility in nitrogen supply during this growth stage. The similar growth observed in the 50% and 100% nitrate treatments suggests that moderate nitrate reduction did not compromise vegetative development, a response that may be especially relevant under saline conditions where growth is already limited by osmotic and ionic constraints.

The maintenance of growth under the 50% nitrate treatment indicates that a moderate reduction in nitrate supply was sufficient to sustain vegetative development even under saline conditions. By contrast, the marked growth inhibition observed under severe nitrate restriction (25%) reveals how nitrogen deficiency can exacerbate the detrimental effects of salinity on plant performance. The simultaneous exposure to salinity and severe nitrogen limitation likely imposed a strong metabolic burden on the plants, further restricting growth and developmental processes (Roșca et al., 2023). The resulting growth inhibition suggests that severe nitrogen limitation further reduces the capacity of tomato plants to cope with saline environments, supporting previous observations that nutritional imbalances aggravate the negative effects of salinity on plant performance and productivity (Al-Gaadi et al., 2024).

Foliar chlorophyll concentration showed a direct relationship with the nitrate dose applied through fertigation. The key role of nitrogen in chlorophyll biosynthesis and chloroplast development explains the higher SPAD values obtained in 100% group plants. (Paponov et al., 2021). Furthermore, the reduction in SPAD values under 25% nitrate likely reflects lower nitrogen allocation to photosynthetic tissues and a consequent decline in pigment synthesis. However, the reduction in chlorophyll content was less pronounced than the decrease in plant growth observed under saline conditions, particularly when comparing plants supplied with 100% and 50% nitrate. This differential response suggests that the photosynthetic apparatus was comparatively more resilient to abiotic stress than developmental processes, allowing plants to maintain chlorophyll levels despite substantial reductions in vegetative growth. Similar observations have been reported by Rivero et al. (2014), who showed that tomato plants exposed to abiotic stress can preserve photosynthetic functionality through adaptive mechanisms that contribute to maintaining cellular homeostasis and stress tolerance (Rivero et al., 2014a). These results indicate that moderate reductions in nitrate supply can be implemented without compromising plant growth or nutritional status, at least during the vegetative stage. While severe nitrate restriction (25%) markedly impaired plant growth and chlorophyll accumulation, plants receiving 50% nitrate maintained physiological performance comparable to that observed under full fertilization. Despite being uncommon in commercial hydroponic setups, the salinity response patterns identified here offer valuable framework for nutrient optimization, laying the groundwork for competitive and environmentally responsible agriculture.

The timing of flowering and fruit set determines the period available for fruit growth and maturation and therefore has a major influence on final yield. In the present study, reproductive phenology was strongly influenced by nitrate availability, whereas the impact of salinity largely depended on plant nutritional status. Plants supplied with 100% and 50% nitrate reached anthesis and fruit set earlier than those subjected to severe nitrate restriction, indicating that adequate nitrogen availability is required not only for floral initiation and inflorescence development but also for supporting the successful transition from anthesis to fruit set. Nitrogen plays a well-established role in meristem activity, protein synthesis and developmental signalling pathways involved in the transition from vegetative to reproductive growth (H. Zhang et al., 2022). Conversely, severe nitrate restriction (25%) likely limited the resources required for floral organ development, resulting in delayed anthesis and slower reproductive progression.

A similar pattern to that observed for vegetative growth and chlorophyll content was found for reproductive development. Plants supplied with 50% nitrate under control conditions exhibited flowering and setting fruit dynamics that were largely comparable to those receiving the full nitrate dose, indicating that a moderate reduction in nitrate supply did not compromise reproductive progression. By contrast, the effects of severe nitrate restriction became increasingly evident as plant development advanced. While differences among treatments were relatively small during the early reproductive stages, the probability of flowering progressively decreased in the 25% nitrate treatment, particularly from the fourth inflorescence onwards, revealing the cumulative effects of prolonged nitrogen limitation. Although salinity is generally associated with delayed flowering and reduced reproductive success in tomato (Roșca et al., 2023), its effects were not uniform throughout the crop cycle. In the present study, the impact of salinity became progressively more evident as successive inflorescences were produced, delaying reproductive progression, especially under severe nitrogen limitation. Nevertheless, plants supplied with 100% nitrate maintained a higher probability of flowering and a faster progression through reproductive stages. The Cox model revealed that plants exposed to salinity and supplied with 100% nitrate often exhibited a greater probability of flowering than non-stressed plants receiving only 25% nitrate. This response reinforces the protective role of adequate nitrate nutrition and suggests that nitrogen availability may partially offset the cumulative constraints imposed by salinity on reproductive development.

A similar synergistic interaction was observed for fruit set, where the combination of salinity and severe nitrate restriction (25%) produced the greatest delays in fructification. This response likely reflects the combined metabolic constraints imposed by salinity and nitrogen deficiency on carbon assimilation and assimilate allocation to reproductive structures. Similar interactions between salt stress and nutritional imbalances have previously been associated with reduced reproductive performance in tomato (Rivero et al., 2014c).

The application of Cox proportional hazard models enabled a robust assessment of reproductive phenology despite the presence of censored observations, providing additional insight into flowering and fruit set dynamics throughout the experimental period. The results demonstrate that nitrate availability is a major determinant of reproductive precocity in tomato. Notably, plants supplied with 50% nitrate-maintained flowering and fruit set dynamics comparable to those observed under full fertilization, supporting the potential of moderate nitrate reduction as a strategy to improve fertilization efficiency without compromising reproductive development, even under saline conditions.

Soluble sugars are among the principal determinants of tomato fruit quality, contributing directly to fruit sweetness and consumer acceptance, and may also function as compatible solutes involved in osmotic adjustment under abiotic stress conditions (P. Zhang et al., 2016). Under control conditions, the reduction in glucose and fructose concentrations observed under severe nitrate restriction suggests that nitrogen availability directly influences carbon assimilation and source-sink relationships during fruit development. Reduced nitrogen supply has frequently been associated with lower photosynthetic capacity and decreased carbohydrate availability for reproductive structures, ultimately limiting sugar accumulation in fruits (Sun et al., 2023). This interpretation is consistent with the lower SPAD values observed under the 25% nitrate treatment, which likely reduced photosynthetic carbon assimilation and consequently limited sugar accumulation in developing fruits. In contrast, salinity completely altered this pattern. Instead of reducing soluble sugar concentrations, fruits from salt-stressed plants maintained elevated glucose and fructose levels regardless of nitrate supply, with a tendency towards higher concentrations under the lowest nitrate dose. This response is consistent with the well documented accumulation of compatible solutes under saline conditions, where soluble sugars contribute to cellular osmotic adjustment and help maintain metabolic activity under stress (Rivero et al., 2014a). Salinity frequently enhances sugar accumulation in tomato fruits, resulting in improved quality attributes such as sweetness and soluble solids content (De Pascale et al., 2001). The increased sugar concentrations observed under low nitrate supply and salinity may also reflect a stress-induced reallocation of carbon towards fruit tissues. Under these conditions, growth limitations imposed by salinity and nitrogen restriction could reduce carbon consumption by vegetative organs, favouring the accumulation of soluble carbohydrates in reproductive sinks. Similar increases in fruit sugar concentration have been reported in hydroponically cultivated tomato subjected to saline irrigation (Al-Gaadi et al., 2024). These results indicate that nitrate deficiency negatively affected sugar accumulation under optimal growing conditions, whereas salinity activated metabolic adjustments that preserved, and even enhanced, fruit carbohydrate content despite reduced nitrogen availability.

Lycopene is the predominant carotenoid in tomato fruit and one of the most important antioxidant compounds contributing to fruit nutritional quality(De Pascale et al., 2001). Under non-stress conditions, the progressive reduction in lycopene concentration observed with decreasing nitrate supply suggests that adequate nitrogen nutrition is required to sustain optimal carotenoid biosynthesis during ripening. Remarkably, salinity abolished the dependence of lycopene accumulation on nitrate availability. Regardless of the nitrogen dose supplied, salt-stressed plants maintained lycopene concentrations that equalled or exceeded those observed in fully fertilized control plants. This response suggests that saline conditions activated metabolic pathways favouring carotenoid accumulation, thereby compensating for the negative effects of nitrogen limitation. Similar responses have been reported previously in tomato, where salinity stimulated carotenoid biosynthesis and antioxidant activity, leading to increased lycopene accumulation despite reductions in vegetative growth (De Pascale et al., 2001). The maintenance of high lycopene concentrations under both 50% and 25% nitrate supply indicates that stress-induced antioxidant responses may become more important than nitrogen availability in regulating carotenoid biosynthesis. Consequently, salinity appears to act as a physiological modulator capable of preserving fruit nutritional quality even under conditions of substantial nitrate reduction. Altogether, these results demonstrate that severe nitrate restriction reduced lycopene accumulation under optimal conditions, whereas salinity maintained elevated carotenoid concentrations independently of nitrate availability, highlighting the remarkable capacity of tomato fruits to preserve antioxidant quality under combined nutritional and saline stress.

Soil salinity and nitrogen availability represent two intersecting environmental factors that continuously shape crop physiology and yield. Our transcriptomic analysis demonstrates that nitrate availability significantly influences both the extent and the magnitude of the genome-wide response to NaCl stress in tomato leaves. Rather than operating through entirely disparate genetic pathways, salinity elicits a substantial core response consisting of 1,314 DEGs consistently regulated across all tested nitrate regimes (25%, 50%, and 100% N). Functional enrichment of this shared transcriptomic core revealed a uniform down-regulation of genes involved in photosystem I and II architecture, light-harvesting processes, carbohydrate metabolism, and cellulose biosynthesis. This consistent repression at the transcript level provides a molecular basis for the physiological growth limitations observed under salinity, where reduced cell expansion and carbon assimilation occur even when visual pigment indices (SPAD) remain transiently stable (Guo et al., 2022; Rivero et al., 2014d). Conversely, the shared up-regulated core was specifically enriched in phosphor relay signal transduction systems and unfolded protein binding (chaperones), reflecting a universal cellular requirement for osmotic signalling and protein conformational protection under ionic stress (Batelli et al., 2024; Su et al., 2021; Zhu, 2016).

Crucially, our data reveal that nitrate availability acts as a quantitative dial governing transcriptional responsiveness to salinity. The highest total number of NaCl-responsive transcripts (6,373 DEGs) and the maximum median response magnitude (|log_2_FC| of 0.545 for up-regulated and 0.731 for downregulated genes) were detected at the intermediate nitrate concentration (50% N), outperforming both the 100% N and 25% N conditions. From a physiological standpoint, plants receiving 50% nitrate sustained vegetative growth, SPAD values, and reproductive phenology (flowering and fruit set kinetics) equivalent to fully fertilized plants. At the transcriptomic level, this intermediate nitrate supply appears to represent a metabolic sweet spot: it avoids severe nutrient starvation while relieving the metabolic burden and osmotic competition associated with excessive nitrate uptake, thereby enabling optimal transcriptional plasticity to deploy adaptive stress pathways (Lopez-Delacalle et al., 2020).

Beyond governing global response magnitude, nitrate availability dynamically altered the transcriptional regulation of a discrete subset of 287 genes exhibiting a statistically significant Nitrate per Salt interaction (adjusted *p*-value *< 0.05*). Functional annotation of this interaction subset pinpointed a specific cluster of 14 genes encoding structural components of the ribosome and translation machinery. Remarkably, these translation-associated transcripts exhibited a directional inversion in response to NaCl depending on nitrogen status: they were uniformly up-regulated under severe nitrogen deficiency (25% N), showed weakened induction at 50% N, and shifted to predominant down-regulation under full nitrogen supply (100% N). In plant systems, translational control plays a paramount role in orchestrating stress survival, often coordinated by central energy sensors such as TARGET OF RAPAMYCIN (TOR) and SNF1-RELATED KINASE 1 (SnRK1) (Margalha et al., 2019; Wu et al., 2024). Under optimal or high nutrient conditions, severe abiotic stress typically prompts SnRK1 activation and ABA-triggered nuclear exit to suppress TOR activity, leading to a global suppression of canonical translation to conserve energy and reallocate resources (Belda-Palazón et al., 2022). This explains the observed repression of ribosomal components at 100% N.

In contrast, under acute nitrogen limitation (25% N), the paradoxical transcriptional upregulation of specific ribosomal subunit transcripts and protein folding chaperones (also reflected in the unique GO enrichment of upregulated processes at 25% N) suggests a cellular emergency response. Under combined severe nitrogen starvation and osmotic shock, plants may upregulate specific ribosomal transcript variants to facilitate localized protein turnover, synthesis of essential protective machinery, or selective translation of stress-responsive mRNAs (Wu et al., 2024). However, as transcript abundance alone does not directly equate to active translation rates or polysome assembly, further translatomic profiling (e.g., Ribo-seq) will be required to confirm whether this transcriptomic shift translates into altered ribosome density or selective mRNA translation in vivo (Wu et al., 2024). Phenotypically, this low-nitrogen emergency strategy coincides with a severe arrest in shoot elongation and delayed reproductive progression, yet results in the preservation and enhancement of fruit soluble sugars (glucose and fructose) and lycopene content. Thus, the transcriptional induction of maintenance machinery under 25% N appears to underpin a survival-oriented metabolic shift, sacrificing vegetative growth to sustain essential cellular homeostasis and preserve fruit sink quality under adverse environmental conditions (Batelli et al., 2024; Rivero et al., 2014d).

In conclusion, our study establishes that nitrate availability is not merely a passive nutritional input, but an active metabolic switch that governs transcriptomic responsiveness, growth-defense trade-offs, and fruit quality in tomato under salinity. By demonstrating that a 50% reduction in nitrate supply maintains vegetative growth, reproductive kinetics, and fruit quality while maximizing transcriptomic plasticity, we identify an optimal fertigation threshold that improves nitrogen-use efficiency without compromising salt stress acclimation. Furthermore, the molecular interplay between nitrogen status and salinity, orchestrated through the SnRK1–TOR metabolic axis and translational reprogramming, explains how tomato plants dynamically sacrifice vegetative growth under severe nutrient limitation to prioritize protective osmolytes and antioxidants in developing fruits. Ultimately, these findings provide a valuable molecular and physiological blueprint for optimizing sustainable fertilization strategies in salt-affected agricultural soils, balancing crop resilience with high-value fruit production.

## ACKNOWLEDGMENTS

We thank the Spanish networks RED2022-134072-T and RED2022-134917-T for critical comments and stimulating discussions on the manuscript.

## AUTHOR CONTRIBUTIONS

J.A.M.-C., S.G.-J., M.C.-P., R.L.-B., and A.C. performed the research. J.A.M.-C., S.G.-J., P.A. and O.L. analysed the data. C.G-S. and Nutricontrol S.L. provided all the sensors and material. J.A.M.-C., S.G.-J., P.A. and O.L wrote the manuscript; and P.A. and O.L. designed the research and supervised the work, O. L. provided financial support. All authors discussed the results and commented on the manuscript.

## SUPPORTING INFORMATION

**Supplementary Figure 1.**
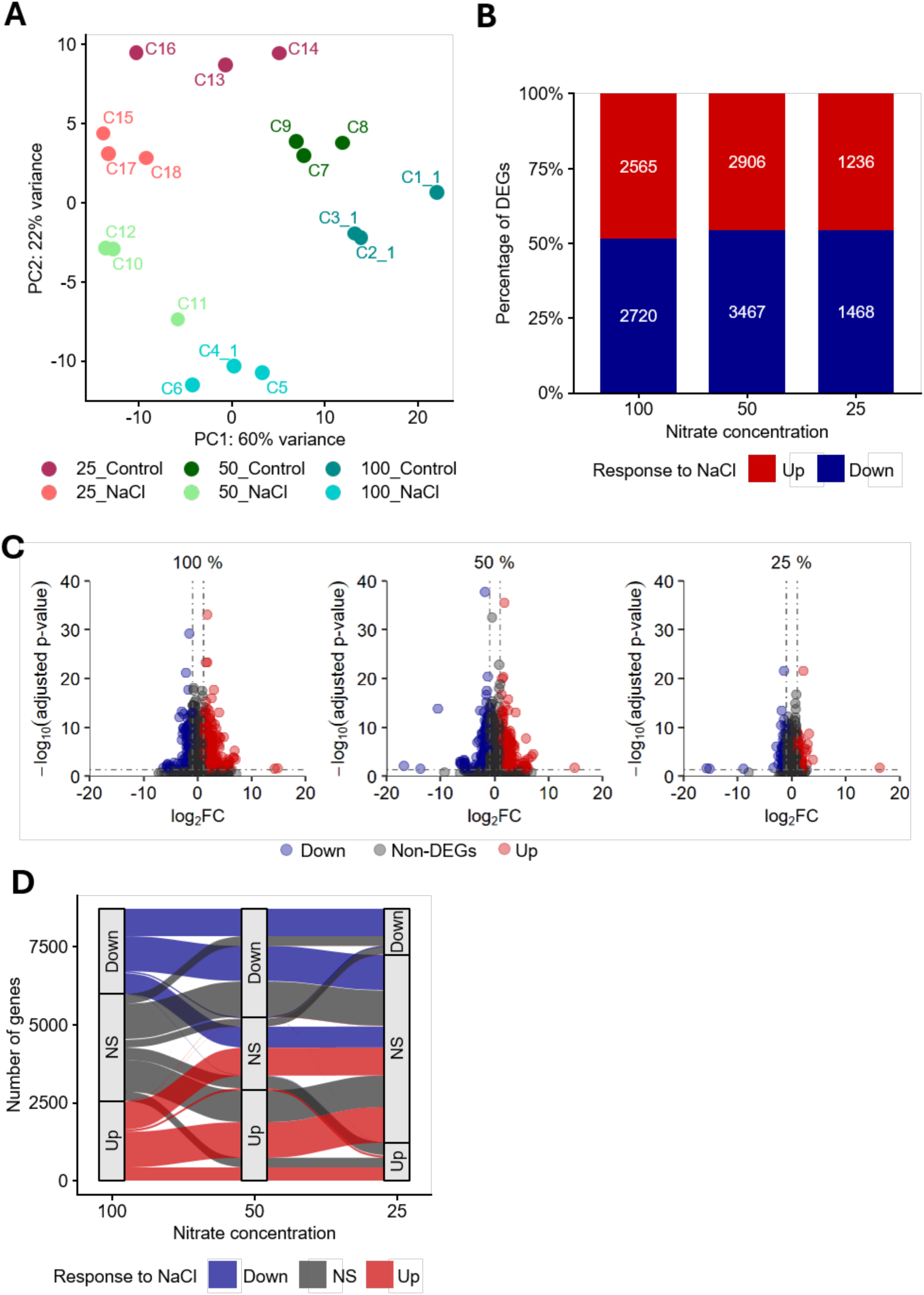
Global transcriptomic responses to NaCl across nitrate concentrations. (A) Principal component analysis of RNA-seq samples across nitrate and NaCl conditions. PCA was performed on variance-stabilizing transformed (VST) count data from all samples. Samples are coloured according to nitrate concentration and NaCl treatment, and individual sample identifiers are shown next to each point. PC1 and PC2 explain 60% and 22% of the total variance, respectively. (B) Distribution of NaCl-responsive differentially expressed genes across nitrate concentrations. Stacked bar plot showing the proportion of up-regulated (red) and down-regulated (blue) genes identified in the NaCl versus Control contrasts at nitrate concentrations of 100, 50, and 25. Numbers within bars indicate the absolute number of genes in each category, while bar heights represent their relative proportions. Genes were considered differentially expressed using an adjusted *p*-value ≤ 0.05. (C) Differential transcriptional response to NaCl at different nitrate concentrations. Volcano plots showing differential gene expression between NaCl-treated and Control samples at nitrate concentrations of 25, 50 and 100. The x-axis represents the log_2_foldchange and the y-axis the −log_10_ adjusted *p*-value. Genes were classified as up-regulated (red) or down-regulated (blue) when the adjusted *p*-value was ≤ 0.05 and the absolute og_2_foldchange was ≥ 1; genes not meeting both criteria are shown in grey. Dashed vertical and horizontal lines indicate the og_2_foldchange (±1) and adjusted *p*-value (0.05) thresholds, respectively. (D) Transitions in NaCl-responsive gene regulation across nitrate concentrations. Alluvial plot showing changes in the regulatory status of genes across the NaCl versus Control contrasts at nitrate concentrations of 100, 50, and 25. Genes were classified as up-regulated, down-regulated, or non-significant (NS) based on an adjusted *p*-value ≤ 0.05. Flows indicate how genes transition between regulatory categories across nitrate concentrations, with red, blue, and grey representing up-regulated, down-regulated, and non-significant genes, respectively.

**Supplementary Figure 2.**
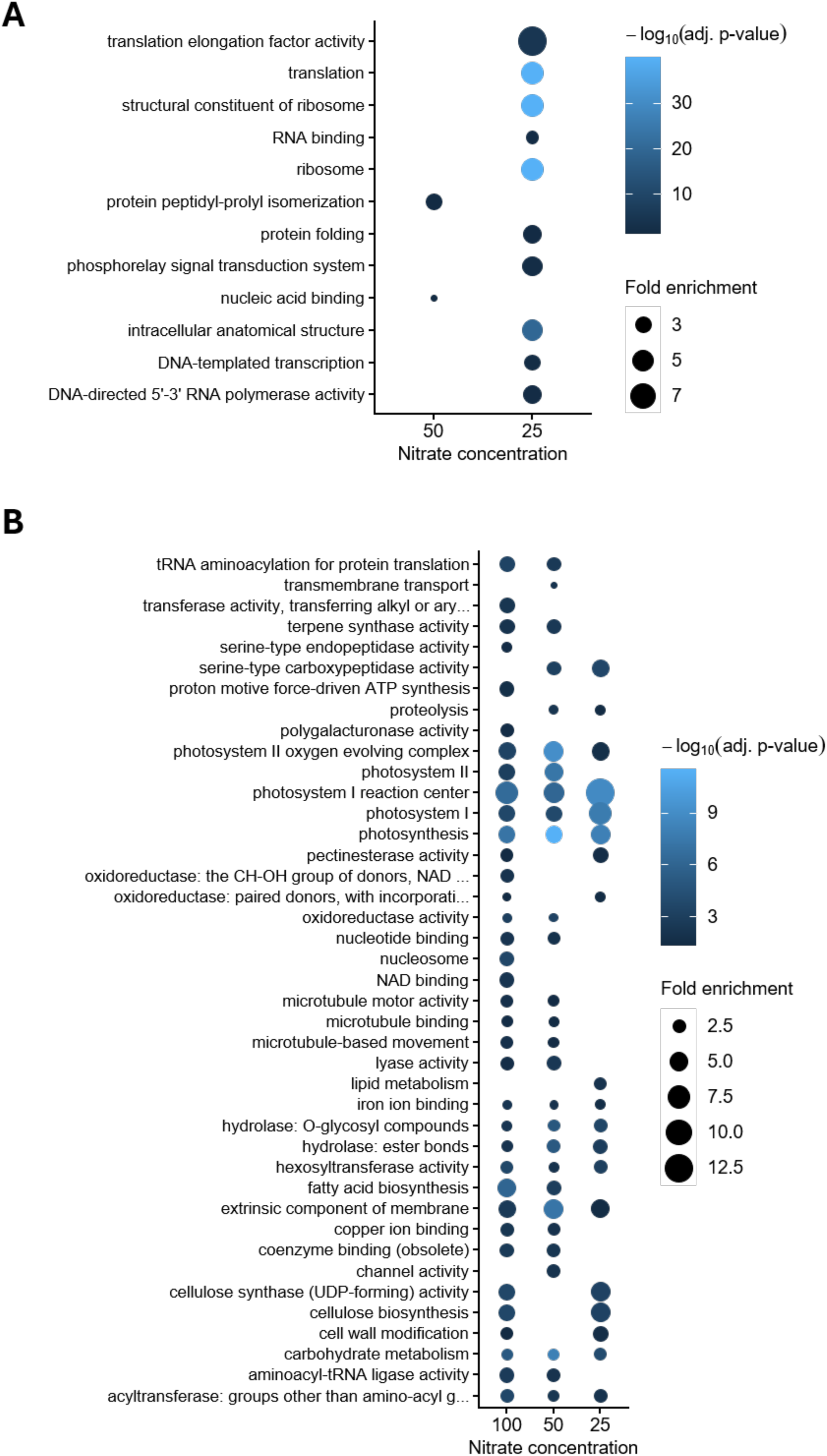
GO enrichment of NaCl-responsive genes at different nitrate concentrations. **(A) GO enrichment of NaCl-induced genes at different nitrate concentrations.** Dot plot showing significantly enriched Gene Ontology terms among genes up-regulated by NaCl at nitrate concentrations of 50 and 25. The x-axis indicates nitrate concentration. Point size represents fold enrichment relative to the GO-annotated background, and point colour represents the adjusted *p*-value. No significantly enriched GO terms were detected among up-regulated genes at nitrate concentration 100. Only terms with adjusted *p*-value ≤ 0.05 are shown. **(B) GO enrichment of NaCl-repressed genes at different nitrate concentrations.** Dot plot showing significantly enriched Gene Ontology terms among genes down-regulated by NaCl at nitrate concentrations of 100, 50, and 25. The x-axis indicates nitrate concentration. Point size represents fold enrichment relative to the GO-annotated background, and point colour represents the adjusted *p*-value. Only terms with adjusted *p*-value ≤ 0.05 are shown.

## REFERENCES

Aishwarya, & Kumar, P. (2024). A Perspective Way to Climate Smart Agriculture. Technological Approaches for Climate Smart Agriculture, 9–24. 10.1007/978-3-031-52708-1_2

Al-Gaadi, K. A., Tola, E. K., Madugundu, R., Zeyada, A. M., Alameen, A. A., Edrris, M. K., Edrees, H. F., & Mahjoop, O. (2024). Response of leaf photosynthesis, chlorophyll content and yield of hydroponic tomatoes to different water salinity levels. PLOS ONE, 19(2), e0293098. 10.1371/JOURNAL.PONE.0293098

Ali, M. Y., Sina, A. A. I., Khandker, S. S., Neesa, L., Tanvir, E. M., Kabir, A., Khalil, M. I., & Gan, S. H. (2020). Nutritional Composition and Bioactive Compounds in Tomatoes and Their Impact on Human Health and Disease: A Review. Foods, 10(1), 45. 10.3390/foods10010045

Andrews, S. R. (2010). FastQC: a quality control tool for high throughput sequence data (0.11.9).

Batelli, G., Ruggiero, A., Esposito, S., Venezia, A., Lupini, A., Nurcato, R., Costa, A., Palombieri, S., Vitiello, A., Mauceri, A., Cammareri, M., Sunseri, F., Grandillo, S., Granell, A., Abenavoli, M. R., & Grillo, S. (2024). Combined salt and low nitrate stress conditions lead to morphophysiological changes and tissue-specific transcriptome reprogramming in tomato. Plant Physiology and Biochemistry, 215(108976), 108976. 10.1016/J.PLAPHY.2024.108976

Belda-Palazón, B., Costa, M., Beeckman, T., Rolland, F., & Baena-González, E. (2022). ABA represses TOR and root meristem activity through nuclear exit of the SnRK1 kinase. Proceedings of the National Academy of Sciences of the United States of America, 119(28), e2204862119. 10.1073/PNAS.2204862119

Cackett, L., Cannistraci, C. V., Meier, S., Ferrandi, P., Pěnčík, A., Gehring, C., Novák, O., Ingle, R. A., & Donaldson, L. (2022). Salt-Specific Gene Expression Reveals Elevated Auxin Levels in Arabidopsis thaliana Plants Grown Under Saline Conditions. Frontiers in Plant Science, 13. 10.3389/FPLS.2022.804716

de Mendiburu, F. (2023). agricolae: Statistical Procedures for Agricultural Research. https://CRAN.R-project.org/package=agricolae

De Pascale, S., Maggio, A., Fogliano, V., Ambrosino, P., & Ritieni, A. (2001). Irrigation with saline water improves carotenoids content and antioxidant activity of tomato. Journal of Horticultural Science and Biotechnology, 76(4), 447–453. 10.1080/14620316.2001.11511392

Dorais, M., Papadopoulos, A. P., & Gosselin, A. (2000). Greenhouse Tomato Fruit Quality. Horticultural Reviews, 239–319. 10.1002/9780470650806.CH5

Fahad, S., Saud, S., Chen, Y., Wu, C., Wang, D., Fahad, S., Saud, S., Chen, Y., Wu, C., & Wang, D. (2021). Abiotic Stress in Plants. Abiotic Stress in Plants. 10.5772/intechopen.91549

Fox, J., Weisberg, S., & Price, B. (2024). Companion to Applied Regression [R package car version 3.1-3]. CRAN: Contributed Packages. 10.32614/CRAN.PACKAGE.CAR

Ghanem, M. E., Van Elteren, J., Albacete, A., Quinet, M., Martínez-Andújar, C., Kinet, J. M., Pérez-Alfocea, F., & Lutts, S. (2009). Impact of salinity on early reproductive physiology of tomato (Solanum lycopersicum) in relation to a heterogeneous distribution of toxic ions in flower organs. Functional Plant Biology, 36(2), 125–136. 10.1071/FP08256

Global status of salt-affected soils. (2024). Global Status of Salt-Affected Soils. 10.4060/CD3044EN

Gohel, D., Moog, S., & Heckmann, M. (2024). Manipulation of Microsoft Word and PowerPoint Documents [R package officer version 0.6.7]. CRAN: Contributed Packages. 10.32614/CRAN.PACKAGE.OFFICER

Guo, M., Wang, X. S., Guo, H. D., Bai, S. Y., Khan, A., Wang, X. M., Gao, Y. M., & Li, J. S. (2022). Tomato salt tolerance mechanisms and their potential applications for fighting salinity: A review. Frontiers in Plant Science, 13(949541). https://doi.org/10.3389/FPLS.2022.949541/FULL

Kassambara, A. (2023a). ggpubr: ggplot2 Based Publication Ready Plots. https://rpkgs.datanovia.com/ggpubr/

Kassambara, A. (2023b). rstatix: Pipe-Friendly Framework for Basic Statistical Tests. https://rpkgs.datanovia.com/rstatix/

Kim, D., Paggi, J. M., Park, C., Bennett, C., & Salzberg, S. L. (2019). Graph-based genome alignment and genotyping with HISAT2 and HISAT-genotype. Nature Biotechnology, 37(8), 907–915. 10.1038/s41587-019-0201-4

Lopez-Delacalle, M., Camejo, D. M., García-Martí, M., Nortes, P. A., Nieves-Cordones, M., Martínez, V., Rubio, F., Mittler, R., & Rivero, R. M. (2020). Using Tomato Recombinant Lines to Improve Plant Tolerance to Stress Combination Through a More Efficient Nitrogen Metabolism. Frontiers in Plant Science, 10(1702). https://doi.org/10.3389/FPLS.2019.01702/TEXT

Love, M. I., Huber, W., & Anders, S. (2014). Moderated estimation of fold change and dispersion for RNA-seq data with DESeq2. Genome Biology, 15(550). 10.1186/s13059-014-0550-8

Margalha, L., Confraria, A., & Baena-González, E. (2019). SnRK1 and TOR: modulating growth-defense trade-offs in plant stress responses. Journal of Experimental Botany, 70(8), 2261–2274. 10.1093/JXB/ERZ066

Millard, S. P. . (2013). EnvStats : an R package for environmental statistics. 291.

Mu, X., & Chen, Y. (2021). The physiological response of photosynthesis to nitrogen deficiency. Plant Physiology and Biochemistry, 158, 76–82. 10.1016/j.plaphy.2020.11.019

Núñez-Ramírez, F., Grijalva-Contreras, R. L., Robles-Contreras, F., Macías-Duarte, R., Escobosa-García, M. I., & Santillano Cázares, J. (2017). Influencia de la fertirrigación nitrogenada en la concentración de nitratos en el extracto celular de peciolo, el rendimiento y la calidad de tomate de invernadero. Revista de La Facultad de Ciencias Agrarias. Universidad Nacional de Cuyo, 49(2), 93–103. https://www.scielo.org.ar/scielo.php?script=sci_arttext&pid=S1853-86652017000200007&lng=es&nrm=iso&tlng=es

Paponov, M., Arakelyan, A., Dobrev, P. I., Verheul, M. J., & Paponov, I. A. (2021). Nitrogen Deficiency and Synergism between Continuous Light and Root Ammonium Supply Modulate Distinct but Overlapping Patterns of Phytohormone Composition in Xylem Sap of Tomato Plants. Plants, 10(3), 573. 10.3390/PLANTS10030573

Park, C. E., Jeong, S. J., Joshi, M., Osborn, T. J., Ho, C. H., Piao, S., Chen, D., Liu, J., Yang, H., Park, H., Kim, B. M., & Feng, S. (2018). Keeping global warming within 1.5 °C constrains emergence of aridification. Nature Climate Change 2017 8:1, 8(1), 70–74. 10.1038/s41558-017-0034-4

Praveen, A., Dubey, S., Singh, S., & Sharma, V. K. (2023). Abiotic stress tolerance in plants: a fascinating action of defense mechanisms. 3 Biotech, 13(3). 10.1007/S13205-023-03519-W

Qin, L., Walk, T. C., Han, P., Chen, L., Zhang, S., Li, Y., Hu, X., Xie, L., Yang, Y., Liu, J., Lu, X., Yu, C., Tian, J., Shaff, J. E., Kochian, L. V., Liao, X., & Liao, H. (2019). Adaption of Roots to Nitrogen Deficiency Revealed by 3D Quantification and Proteomic Analysis. Plant Physiology, 179(1), 329–347. 10.1104/pp.18.00716

Quinet, M., Angosto, T., Yuste-Lisbona, F. J., Blanchard-Gros, R., Bigot, S., Martinez, J. P., & Lutts, S. (2019). Tomato Fruit Development and Metabolism. Frontiers in Plant Science, 10. 10.3389/fpls.2019.01554

R Core Team. (2023). R: A Language and Environment for Statistical Computing. https://www.R-project.org/

Rivero, R. M., Mestre, T. C., Mittler, R., Rubio, F., Garcia-Sanchez, F., & Martinez, V. (2014a). The combined effect of salinity and heat reveals a specific physiological, biochemical and molecular response in tomato plants. Plant, Cell & Environment, 37(5), 1059–1073. 10.1111/pce.12199

Rivero, R. M., Mestre, T. C., Mittler, R., Rubio, F., Garcia-Sanchez, F., & Martinez, V. (2014b). The combined effect of salinity and heat reveals a specific physiological, biochemical and molecular response in tomato plants. Plant, Cell and Environment, 37(5), 1059–1073. https://doi.org/10.1111/PCE.12199;ISSUE:ISSUE:DOI

Rivero, R. M., Mestre, T. C., Mittler, R., Rubio, F., Garcia-Sanchez, F., & Martinez, V. (2014c). The combined effect of salinity and heat reveals a specific physiological, biochemical and molecular response in tomato plants. Plant, Cell and Environment, 37(5), 1059–1073. 10.1111/PCE.12199

Rivero, R. M., Mestre, T. C., Mittler, R., Rubio, F., Garcia-Sanchez, F., & Martinez, V. (2014d). The combined effect of salinity and heat reveals a specific physiological, biochemical and molecular response in tomato plants. Plant, Cell & Environment, 37(5), 1059–1073. https://doi.org/10.1111/PCE.12199

Roșca, M., Mihalache, G., & Stoleru, V. (2023). Tomato responses to salinity stress: From morphological traits to genetic changes. Frontiers in Plant Science, 14, 1118383. https://doi.org/10.3389/FPLS.2023.1118383/FULL

Su, X., Wang, B., Geng, X., Du, Y., Yang, Q., Liang, B., Meng, G., Gao, Q., Yang, W., Zhu, Y., & Lin, T. (2021). A high-continuity and annotated tomato reference genome. BMC Genomics, 22(1), 1–12. https://doi.org/10.1186/S12864-021-08212-X/FIGURES/5

Sun, J., Jin, L., Li, R., Meng, X., Jin, N., Wang, S., Xu, Z., Liu, Z., Lyu, J., & Yu, J. (2023). Effects of Different Forms and Proportions of Nitrogen on the Growth, Photosynthetic Characteristics, and Carbon and Nitrogen Metabolism in Tomato. Plants, 12(24), 4175. 10.3390/PLANTS12244175

Tang, H., Zhang, X., Gong, B., Yan, Y., & Shi, Q. (2020). Proteomics and metabolomics analysis of tomato fruit at different maturity stages and under salt treatment. Food Chemistry, 311. 10.1016/j.foodchem.2019.126009

Wickham, H. (2016). ggplot2: Elegant Graphics for Data Analysis. Springer-Verlag New York. https://ggplot2.tidyverse.org

World Food and Agriculture – Statistical Yearbook 2024. (2024). World Food and Agriculture – Statistical Yearbook 2024. 10.4060/CD2971EN

Wu, H. Y. L., Jen, J., & Hsu, P. Y. (2024). What, where, and how: Regulation of translation and the translational landscape in plants. The Plant Cell, 36(5), 1540– 1564. 10.1093/PLCELL/KOAD197

Xu, G., Fan, X., & Miller, A. J. (2012). Plant nitrogen assimilation and use efficiency. Annual Review of Plant Biology, 63, 153–182. 10.1146/ANNUREV-ARPLANT-042811-105532

Yu, G., Wang, L. G., Han, Y., & He, Q. Y. (2012). clusterProfiler: an R Package for Comparing Biological Themes Among Gene Clusters. OMICS : A Journal of Integrative Biology, 16(5), 284–287. 10.1089/OMI.2011.0118

Zhang, H., Zhu, J., Gong, Z., & Zhu, J. K. (2022). Abiotic stress responses in plants. Nature Reviews. Genetics, 23(2), 104–119. 10.1038/S41576-021-00413-0

Zhang, P., Senge, M., & Dai, Y. (2016). EFFECTS OF SALINITY STRESS ON GROWTH, YIELD, FRUIT QUALITY AND WATER USE EFFICIENCY OF TOMATO UNDER HYDROPONICS SYSTEM. Reviews in Agricultural Science, 4(0), 46–55. 10.7831/RAS.4.46

Zhu, J. K. (2016). Abiotic Stress Signaling and Responses in Plants. Cell, 167(2), 313–324. 10.1016/j.cell.2016.08.029

